# Adaptation of *Enterococcus faecalis* to intestinal mucus revealed by a human colonic organoid model

**DOI:** 10.1101/2025.08.20.670035

**Authors:** Sofya Mikhaleva, Po-Long Hsiao, Amanzhol Kurmashev, Caleb M. Anderson, Cristina Colomer-Winter, Julia A. Boos, Pei Yi Choo, Julia L.E. Willett, Andreas Hierlemann, Kimberly A. Kline, Alexandre Persat

## Abstract

The human gastrointestinal tract hosts a diverse population of microorganisms that have a significant impact on host health. Among this population, *Enterococcus faecalis* (*Ef*) represents a common member of intestinal microbiota colonizing humans early in life, but which can also opportunistically infect its host. Despite its importance in human health, investigations of its physiological adaptation to the mucosal environment remain limited. Building on recent advances in tissue engineering, we here leverage human colonic organoids (colonoids) to investigate the *Ef*’s mechanisms of mucosal surface colonization across space and time. Using high-resolution microscopy, we visualized *Ef* growth within the natively formed colonic mucus layer in colonoids. Leveraging a custom perfusion chamber, we tracked *Ef* growth within the mucus of live colonoids over time under flow, which revealed specific colonization strategies, including biofilm-like microcolony formation. To identify *Ef* fitness determinants in this niche, we implemented transposon insertion sequencing (Tn-seq) in the natively formed mucus of live colonoids. This approach revealed a large fitness rearrangement compared to typical liquid culture, mainly involving metabolic activity and regulatory response during mucosal colonization, as well as factors that may contribute to colony formation at the mucosal surface. Altogether, our results show important physiological and biophysical adaptation of *Ef* to the mucosal surface that are not captured by *in vitro* conditions and that cannot be revealed *in vivo* at high resolution.

## Introduction

The human gastrointestinal tract (GI) hosts communities of bacterial species collectively known as the gut microbiota. Many members play key roles in digesting complex dietary fibers, strengthening gut barrier integrity and shaping the host immune system (1, 2). Despite these important functions, the mechanisms by which any given bacterial species stably colonizes the gut remain poorly characterized, in part due to our limited ability to monitor host colonization in space and time at high resolution. The large intestine hosts a particularly abundant bacterial population, physically separated from the colonic epithelium by a thick mucus layer. Mucus is a hydrogel secreted by dedicated goblet cells, predominantly built from the crosslinked chains of heavily glycosylated MUC2 mucin proteins. While the main function of mucus is to physically repel microbes from host epithelial cells, it also acts as a nutrient source to some bacterial species. Many species can degrade mucus and release sugars to nourish the rest of the microbiota (3, 4). As mucus is a complex, heterogeneous material, meticulously investigating its dual function remains complex in animal models, which face limitations in live dynamic visualizations. Conversely, *in vitro* models building on purified mucus often fail to recapitulate its physical integrity upon purification. Consequently, our understanding of colonization and stability of pathogens and commensal microbiota at the mucosal surface is still limited (5–7).

*Enterococcus faecalis* (*Ef*), a Gram-positive non-motile facultative aerobe, is a core member of the human intestinal microbiota (8, 9). As a highly adaptable commensal and one of the earliest GI tract colonizers (10–12), *Ef* supports intestinal physiology by regulating pH, facilitating metabolism, and producing vitamins (13). However, under certain perturbations, including antibiotic-induced dysbiosis or inflammatory bowel disease, *Ef* can become an opportunistic pathogen breaching the epithelial barrier, causing serious infections like bacteremia and endocarditis (14–18). These pathogenic traits are linked to exotoxin production, antibiotic or bile acids tolerance and biofilm formation (19, 20). *Ef* may also employ physical strategies to persist in the host. It specifically grows into multicellular mucus-associated structures that resemble biofilms, thus likely offering protection from flow-induced clearance (5). Despite *Ef*’s role in early GI tract colonization and pathogenicity, most existing knowledge of *Ef* biofilms derives from studies conducted in abiotic environments. These studies have revealed that sortase-regulated adhesins mediate initial surface attachment (21), while secretion of polysaccharides (22, 23) and extracellular DNA (24) promotes biofilm maturation and cohesion. Commitment to form biofilms by any species has a strong impact on bacterial interaction with its environments, for example by restricting nutrient access, which can induce metabolic reprogramming (25, 26). The way in which *Ef* and other microbiota species adapt their physiology to form biofilms at mucosal surfaces remains largely unexplored (5, 27).

To elucidate the mechanisms of bacterial colonization in a more physiologically relevant context, recent advances in tissue and organoid engineering offer a promising alternative. Human-derived organoids bypass limitations of animal experiments, which do not always reflect human physiology. In addition, careful organoid engineering enables high-resolution microscopy and functional genomics integration. For example, we have previously demonstrated that human airway organoids can highlight *Pseudomonas aeruginosa* mechanisms of biofilm formation and pathogenic adaptation during growth and antibiotic treatment (28, 29). Intestinal organoids have also been employed to investigate enterohemorrhagic *Escherichia coli* (30)*, Salmonella* (31)*, Clostridioides difficile* (32)*, Shigella flexneri* infections of the GI tract (33), as well as *Vibrio cholerae* biofilm formation on immune cells after breaking intestinal barriers (34).

To investigate *Ef* adaptation strategies to the gut mucosal barrier in space and time, we developed a framework combining mucus-producing tissues from human colonic organoids (colonoids), live imaging and a functional genomic screen. Fluorescent microscopy revealed that *Ef* grows in mucus-rich regions by forming dense microcolonies, akin to those observed in mouse and human intestines (12, 35). Using transposon sequencing (Tn-seq), we identified key factors involved in *Ef*’s metabolic and physical adaptation to the gut mucosal environment. While several known biofilm-associated factors in our Tn-seq results were dispensable during early colonization, the glycosyltransferase BgsB emerged as critical for *Ef*’s initial adaptation to mucus. We propose that this combined platform of human colonoids, live imaging, and functional genomics offers a broadly applicable tool to study adaptation and pathogenicity in intestinal pathogens, commensals and microbial communities.

## Results

### *Ef* colonizes colonoid mucus

To investigate how *Ef* thrives in a human mucosal environment, we use patient-derived healthy descending colon organoids. Widely used for developmental studies, organoids typically grow in a cystic shape, wherein the epithelial lumen faces inwards (36). Colonization and infection thus require microinjection to access the mucosal surface, thereby undermining organoid integrity (37). To overcome this limitation, we leveraged the open apical configuration of organoids grown on a 2D membrane (Transwell) (38, 39), forming a polarized epithelium with the apical side facing upwards. This allows us to directly access the mucosal surface for microbial inoculation without perturbing the tissue (**Figure 1A**).

**Figure 1.**
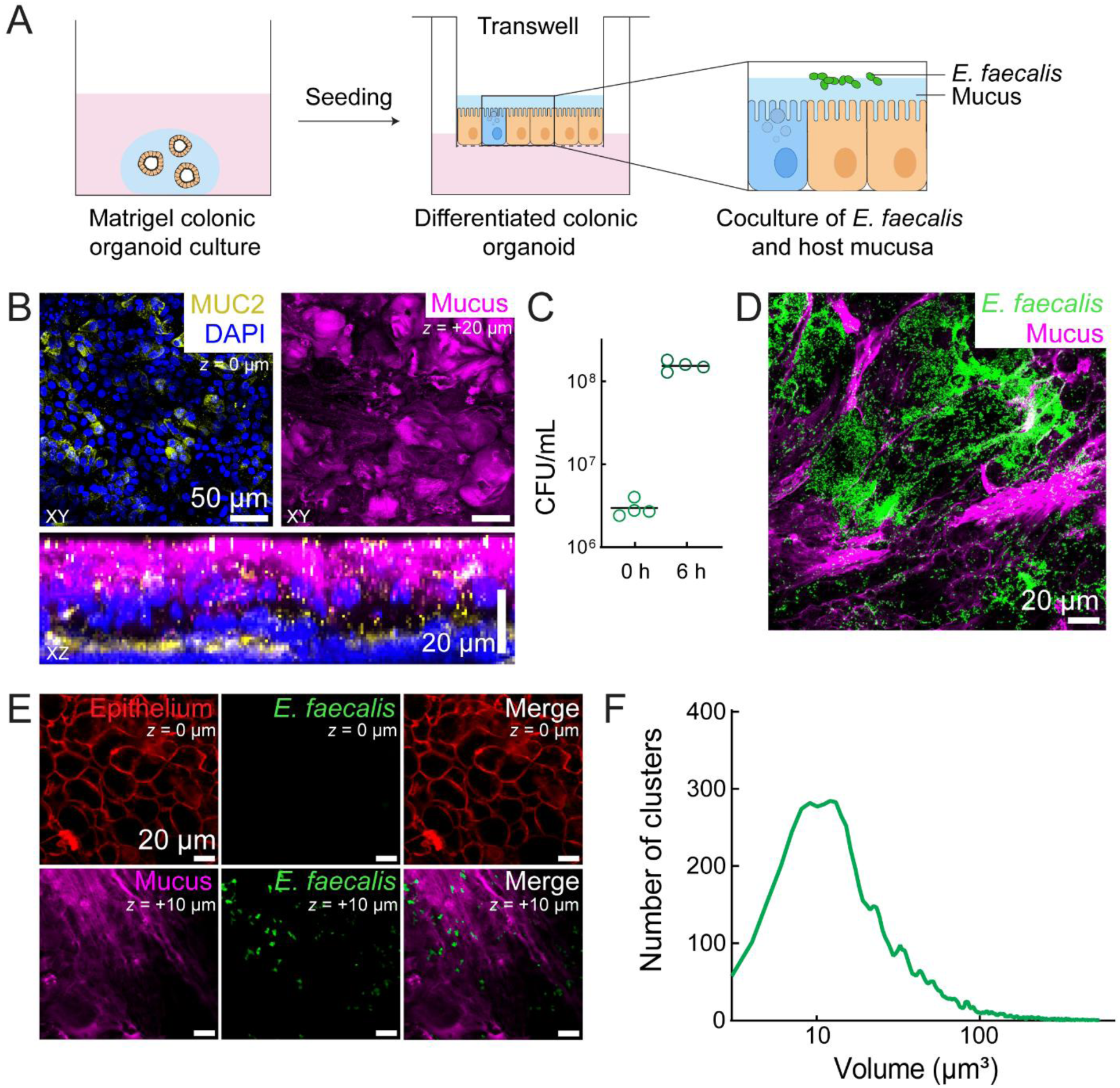
Colon epithelial monolayers with histological signatures for studying *Enterococcus faecalis* mucosal colonization. (A) Schematic depicting the experimental setup. Human colon organoids grown in Matrigel were split into single cells and seeded on top of collagen-coated Transwell membranes. Once the epithelial monolayer formed, the cells were differentiated at the air-liquid interface. Transwells were used on day 7 after seeding. (B) Immunostaining images of differentiated organoid monolayers with MUC2 protein (yellow), nuclei stained with DAPI (blue), and mucus labeled using Jacalin-biotin and Streptavidin-Cy5 (magenta). (C) Colony forming units (CFU/ml) analysis of *Ef* growth in mucus 6 hours after inoculation. Each point is a biological replicate. The horizontal black lines mark the mean. (D) Maximum intensity projection of Jacalin-labeled mucus (magenta) and *Ef* WT expressing pDasherGFP (green). (E) Colonoid epithelium labeled with CellMask-DeepRed (red) is away from the *Ef* WT expressing pDasherGFP (green), which is in proximity with Jacalin-labeled mucus (magenta). (F) *Ef* colony volume analysis from confocal images.

Differentiated colonoids with mucus-secreting goblet cells were shown by immunostaining with anti-MUC2 antibody (**Figure 1B**). To produce an *in vivo*-like mucus quantity, we enhanced mucin production by supplementing the culture with vasoactive intestinal peptide (VIP) (39), resulting in its visible accumulation. To confirm the presence of gel-forming mucins at the epithelial surface, fixed colonoids were labeled with a fluorescent conjugate of the lectin Jacalin, which binds human colonic MUC2 O-glycans (40, 41). Despite extensive washing during fixation and staining steps, which erode mucosal content, Jacalin-staining revealed a ∼ 10 µm thick extracellular mucus layer (**Figure 1B**), mimicking the native colonic environment. We therefore used this platform to interrogate *Ef’s* physiology in intestinal mucus.

We first tested the ability of *Ef* to grow at the mucosal surface of 2D colonoids. In this configuration, organoids are exposed to media through the porous membrane at the basal side, while we maintained an air-liquid interface at the apical side, which promotes mucus accumulation. This setup allowed us to assess *Ef’s* adaptation to the mucosal surface, where bacteria experience mucus without exogenous nutrients. To prevent nutrient carryover from prior cultures, *Ef* were thoroughly washed and inoculated in a salt solution in small volumes that preserved the air-liquid interface. To monitor bacterial growth, we quantified colony-forming units (CFU) per Transwell and compared them to the initial inoculum. After 6 h, the number of viable *Ef* cells increased approximately 50-fold, indicating favorable growth conditions (**Figure 1C**). This corresponds to a doubling time of ∼1 h, which is slower than in the rich medium (42), suggesting *Ef* experiences nutrient-limited conditions. Altogether, these experiments indicate that *Ef* potentially utilizes host-derived mucin glycans and amino acids to grow at the mucosal surface.

Next, we investigated the patterns by which *Ef* populations colonize the mucosal surface of colonoids. To visualize the spatial organization of *Ef* around mucus, we leveraged the optical accessibility of human colonoid tissues. We colonized colonoids with an *Ef* strain constitutively expressing GFP (43), and the mucus was stained with Jacalin-biotin and Streptavidin-Cy5 before imaging the tissues with 3D confocal spinning disk microscopy (**Figure 1D**). We could observe that *Ef* preferentially formed microcolonies in mucus-rich areas, where it remained separated from epithelial cells (**Figure 1E**). Cluster size analysis revealed a mean volume of ∼28 µm³ (**Figure 1F**). Assuming the cluster grew from a single 0.5 µm³ bacterial cell (1 µm-diameter sphere), volume expansion approached 50-fold, consistent with CFU measurements (**Figure 1C**). These observations together suggest that *Ef* not only survives but proliferates at the colonoid mucosal surface without external nutrients, forming large microcolonies reminiscent of biofilms. We therefore hypothesize that *Ef* exploits mucus-derived nutrients to sustain growth and establishes a stable, host-adapted lifestyle at the mucosal surface.

### *Ef* forms multicellular clusters embedded in mucus

To investigate the morphogenesis of *Ef* biofilms at the mucosal surface, we performed time-lapse microscopy under flow that replicates the conditions experienced in the GI tract. To enable the application of controlled flow, we grew colonoids at the bottom surface of the Transwell membrane rather than the top, in a so-called inverted configuration. We then used a custom microfabricated microscopy adaptor that precisely positions the tissue in close proximity to the coverslip surface for high-resolution imaging (44). The device also incorporates channels with inlets and outlets, enabling steady fluid perfusion (Figure 2A). In addition to replicating intestinal flow, this setup also removes detached bacterial biomass, enabling accurate tracking of mucus-bound populations. We pre-labeled mucus with Jacalin-biotin and Streptavidin-Cy5 before inoculating GFP-expressing *Ef* directly onto the mucus. We used a flow intensity on the same order as the one experienced in the large intestine (45).

**Figure 2.**
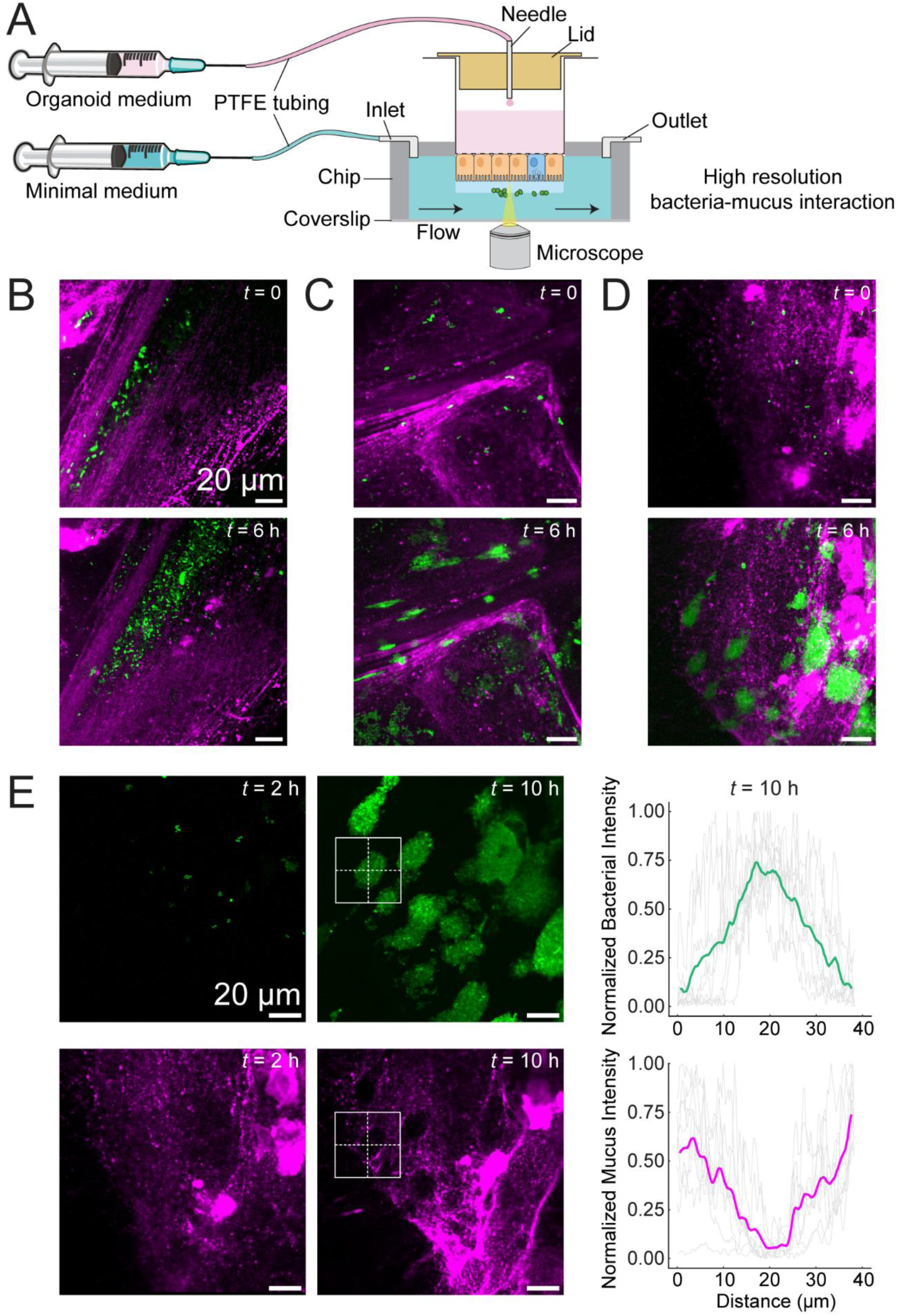
*Ef* grows in cavities surrounded by mucus. (A) Schematic depicting the experimental setup using a Transwell-adapted microfluidic device for high-resolution confocal imaging. (B) *Ef* colony growing between thick mucus fibers and not forming biofilm-like colonies. (C) *Ef* colony growing inside the mucus but in areas with low mucus labeling. (D) *Ef* colony growing and slowly pushing Jacalin-labeled mucus away upon expansion. (E) Images of the bacterial channel (green) and mucus channel (magenta) of the timelapse confocal imaging of *Ef* in mucus under flow. Graphs represent the average of quantification of the sum of the vertical and horizontal profile measurements (dashed lines within the cropped region) of each colony analyzed in the bacterial channel (upper right) and mucus channel (lower right) (n = 9).

This setup allowed us to perform live microscopic tracking of bacterial growth at the mucosal surface. We first examined *Ef* growth over time at defined regions of the mucosal surface. *Ef* grew into groups of different densities, consistently localized away from epithelial cells but always near mucus-rich regions. Visualizing *Ef* at single-cell resolution revealed heterogeneity in growth behaviors. One subpopulation grew into low-density communities within large spaces formed between mucus strands (Figure 2B). Another subpopulation grew into densely packed clusters of contiguous *Ef* cells confined within narrow mucus-lined trenches (Figure 2C). Additionally, some bacterial clusters were embedded within mucus-filled spherical cavities (Figure 2D). Overall, we observed limited overlap between *Ef* and Jacalin-labeled mucus. Fluorescence intensity quantification confirmed that most clusters grew in areas of low mucus fluorescence, surrounded by a strong mucus signal (Figure 2E), suggesting that the elastic gel-like structure of mucus supports and actively shapes *Ef* growth. Together, these findings show that under flow, *Ef* preferentially colonizes and grows within Jacalin-labeled mucus cavities, rather than at the surface.

### Transposon sequencing in colonoids reveals *Ef* mucus adaptation strategies

Our imaging data indicate that *Ef* rapidly adapts to the mucosal surface, growing using host-derived nutrients. Its slower growth rate and biofilm formation suggest underlying metabolic reprogramming and specialized phenotypes required for mucus adaptation. We hypothesized that these adaptive strategies differ significantly from those used in rich medium cultures. To identify genetic factors contributing to growth on mucus, we conducted a high-throughput functional genomic screen in colonoids. We performed transposon sequencing (Tn-seq) using a *Himar*-based transposon library of the OG1RF strain with 1,926 insertions of 2,651 total annotated open reading frames (46). We anticipated that mutants in genes important for adaptation to mucus would show reduced fitness.

We cultured the Tn library and compared its fitness in the colonic mucus conditions to growth in the standard brain-heart infusion (BHI) medium, which was used to grow the library prior to inoculation. To control for interference from organoid medium leakage through the epithelium, we sequenced a second reference condition with the library growing in the basal medium used in colonoid tissue culture (Figure 3A). Incubation times were limited to 6 division cycles (**Figure S1A**), during which *Ef* forms large colonies in the mucus layer but does not cause any large-scale tissue damage as observed by propidium iodide staining (**Figure S1B**). Under these conditions, we expected that selection would primarily reflect the inability of mutants to grow in mucus, rather than effects from epithelial damage.

**Figure 3.**
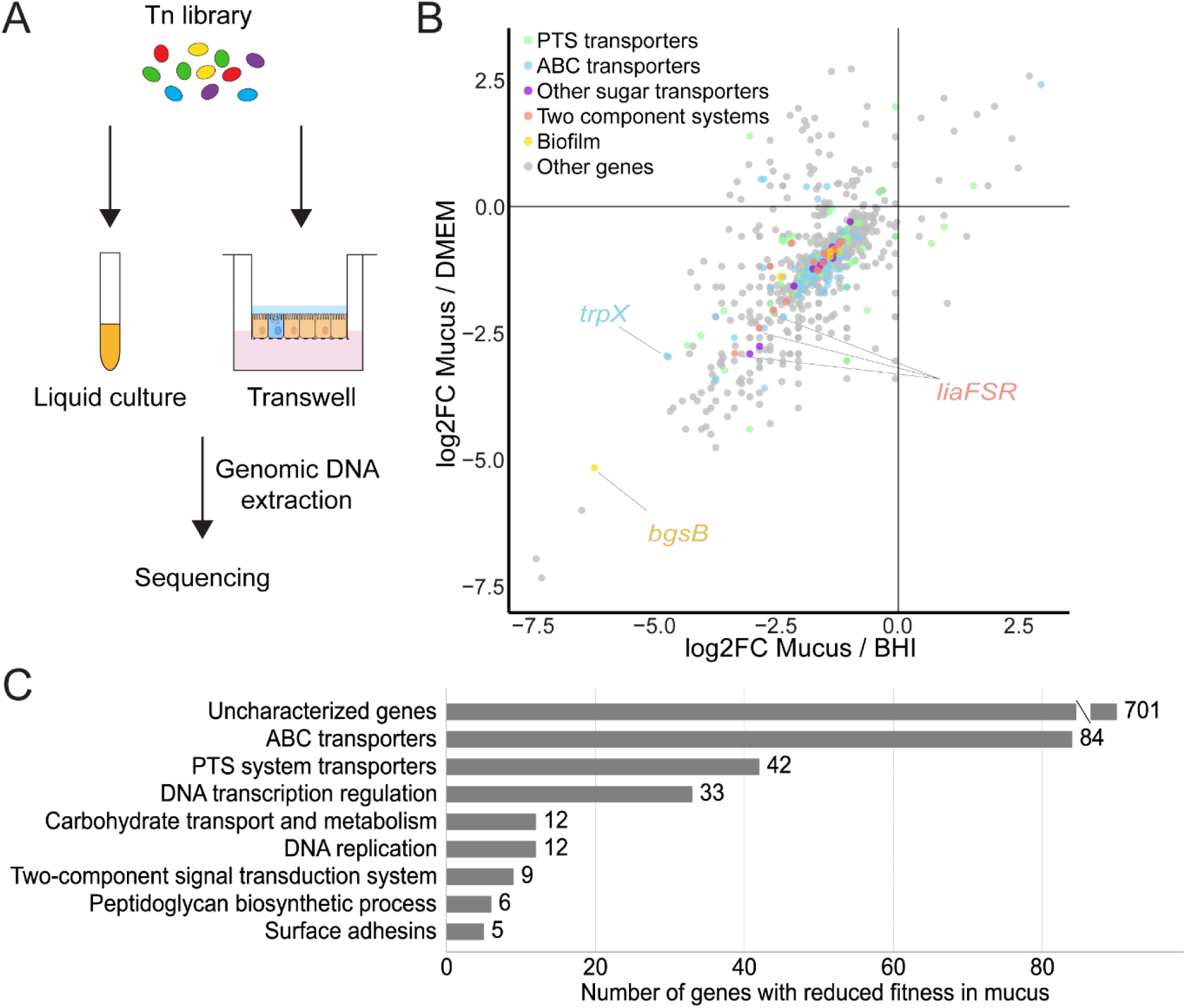
Tn-seq in colonoids reveals mechanisms of adaptation to the mucosal surface. (A) Scheme depicting the experimental setup for comparing *Ef* growth in colonoids *vs* liquid media. (B) Scatter plot representing the distribution of transposon mutants in mucus with highlighted carbohydrate transporters in blue, biofilm-related genes in yellow, and two-component signal transduction adaptation system genes in red. Genes with the largest fold change in reduced fitness in these categories were highlighted. (C) Biological processes found in a gene ontology analysis of the Tn-seq results (KEGG Pathway).

Analysis of the Tn-seq identified 569 genes with reduced fitness (log2FC < −2) in mucus compared to growth in BHI, many of which were annotated as hypothetical. Among genes of known function, we identified genes responsible for metabolism that include ABC and PTS system transporters and carbohydrate transporters, indicating adaptation to distinct nutrient conditions and consistent with the reliance on host-derived glycans. Furthermore, mutations in genes associated with environmental adaptation, like two-component signal transduction systems, suggest that *Ef* modulates its physiology in response to specific inputs at the mucosal surface. Finally, mutations in biofilm factors such as peptidoglycan biosynthesis and surface adhesins indicate that the biofilm lifestyle may promote fitness in the mucosal environment (Figure 3B and C).

### *Ef* metabolically adapts to mucus

Tn-seq in mucus identified metabolic and two-component system genes to be important in mucosal adaptation (Figure 3C). Mucins, the primary glycoproteins in mucus, are heavily glycosylated and contain a variety of sugar residues in their O- and N-glycans, including mannose, fucose, galactose, N-acetylglucosamine (GlcNAc), N-acetylgalactosamine (GalNAc) and sialic acids (3, 4). Our Tn-seq screen showed strong fitness defects for mutations in genes involved in tryptophan and methionine biosynthesis, as well as an operon of galactose/GalNAc phosphotransferase transporters (**Figure S2A**). We validated this observation and observed that mutations in *trpX* and *metQ* genes resulted in ∼30% reduced growth rate compared to WT (**Figure S2B**).

Additionally, we identified two previously uncharacterized metabolic genes of the ABC-family sugar transporters that are important for *Ef* growth on mucus (**Figure S2A**). We found that mutations in *OG1RF_10303* and *OG1RF_12205* result in a significant fitness reduction in mucus (**Figure S2B**). Consistent with our Tn-seq results, we found that these mutants had no growth defects in rich medium. To identify the mechanisms by which they help *Ef* metabolically adapt at the mucosal surface, we screened the growth of these mutants in a minimal medium supplemented with sugar residues that can be released from mucus. We found that a mutant in *OG1RF_11614* results in a strong fitness defect in GalNAc, while the growth of *OG1RF_10303* and *OF1RF_12205* mutants was unaffected by the sugars we could screen (**Figure S2C**). These data confirm that *Ef* deploys a metabolic activity compatible with nutrient availability at the mucosal surface.

Furthermore, the functional genomics screen identified 3 two-component sensory systems (the *Ef* OG1RF genome encodes 16) that may play a role in mucosal adaptation (**Figure S3A**). Mutations in the *croRS*, *liaFSR,* and *bsrRS* two-component systems decreased fitness. We validated the reduced fitness of *bsrRS* and *croRS* mutants in colonoids by CFU analysis, confirming that *Ef* rapidly senses specific signals to adapt to the intestinal mucosal environment (**Figure S3B**). Which signals activate these systems at the mucosal surface remains however unclear.

### BgsB is essential to multicellular clusters formation in mucus

Given the formation of large biofilm-like colonies in mucus, we investigated whether interfering with this process could impact *Ef* fitness. We mined the Tn-seq data for genes linked to biofilm formation on abiotic surfaces and identified two with significant fitness defects when grown in mucus (Figure 4A). One was *bph*, a biofilm-forming factor whose deletion decreases the expression of the *fsr* locus, gelatinase (*gelE*), and multiple cell surface WxL domain proteins, all of which contribute to *in vitro* biofilm formation and surface attachment (47, 48). Another candidate was *bgsB*, a glycosyltransferase required for diglucosyldiacylglycerol synthesis and biofilm formation (49).

**Figure 4.**
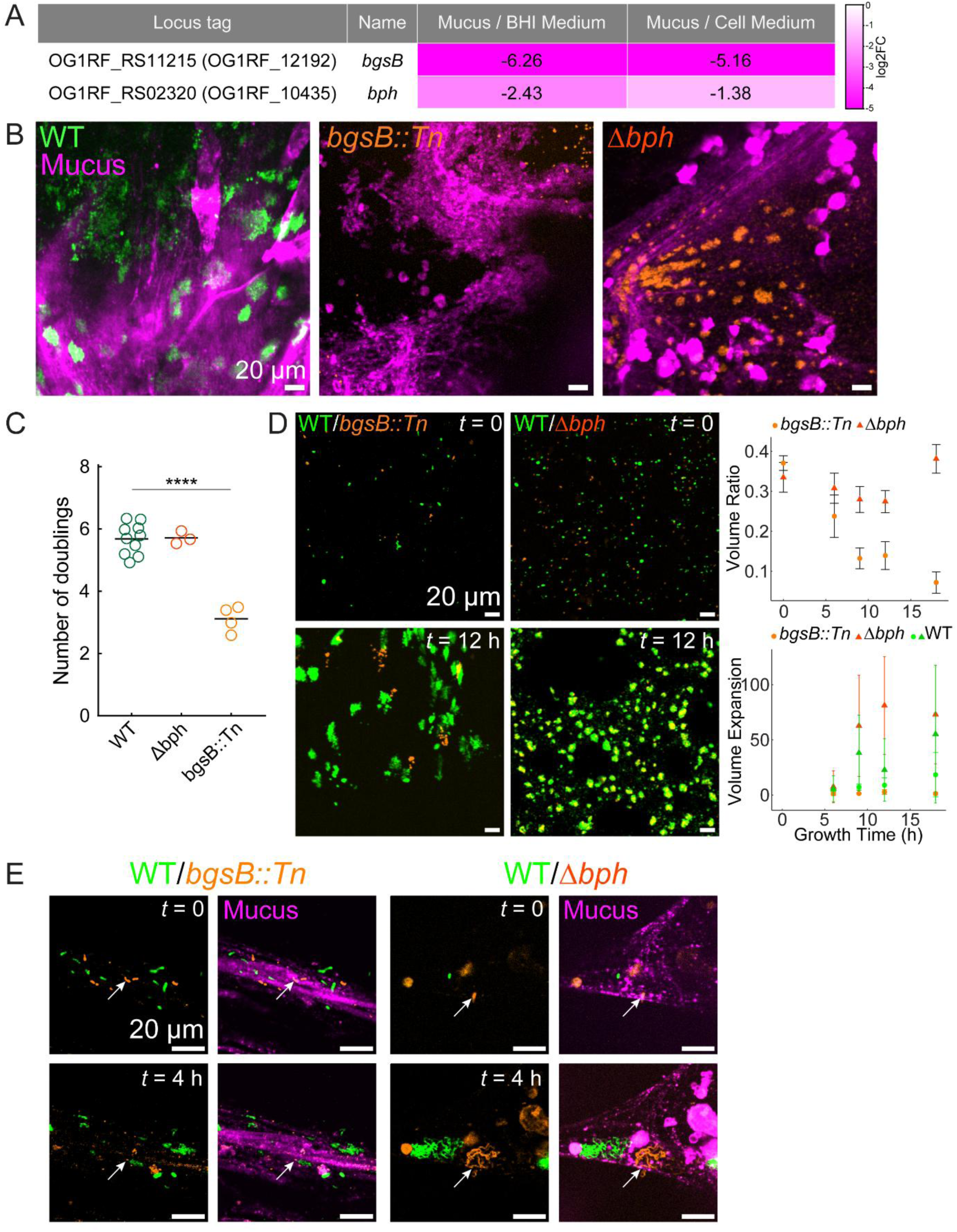
Glycosyltransferase BgsB in *E. faecalis* is essential for growth in colonic mucus. (A) Table of log2FC values calculated from the Tn-seq experiment for the glycosyltransferase *bgsB* and a biofilm-forming factor *bph*. (B) *Ef* colonies in colonic mucus at 9 hours of growth.

We confirmed that the *bgsB::Tn* mutant exhibited significantly reduced fitness in mucus compared to WT (Figure 4B and C). By contrast, CFU did not confirm a fitness defect for the *bph* deletion mutant. Growing in minimal medium conditions, the *Δbph* mutant showed a delayed logarithmic growth when mannose was the only carbon source compared to WT (**Figure S4**). To assess how *bgsB* impacts *Ef* mucosal colonization, we returned to imaging. Confocal microscopy showed that the *bgsB::Tn* mutant failed to form biofilm-like microcolonies, while the *Δbph* mutant formed WT-like colonies both in monocultures and co-culture with WT at a 1:1 ratio (Figure 4B-D). High-resolution imaging of *Ef* growth with mucus fibers using the Transwell-based microfluidic platform confirmed the growth defect of the *bgsB::Tn* mutant and showed similar growth for WT and the *Δbph* mutant (Figure 4E).

Mucus was labeled using Jacalin-biotin and Streptavidin-Cy5, the wild-type *Ef* was fluorescently expressing pDasher-GFP and mutants of *bgsB* and *bph* expressing tdTomato. (C) Colony forming units (CFU/ml) quantification of the deletion mutants grown in mucus in comparison with the wild-type *Ef* at 6 hours of growth. The horizontal black lines mark the mean. (D) Competition between WT (green) and mutant samples (orange). Volume ratio is calculated as a fraction of the mutant signal occupying the total (mutant + WT) volume in the image. Volume expansion is calculated as a ratio to the initial volume. Mean and standard deviation are shown. (E) The same mixed samples were grown in mucus in a flow chip, under the flow of 5 µl/min of minimal medium. Arrows indicate the same place at the start and the end of the timelapse. Statistics in (C), Welch unpaired t-test (two-sided) (****P < 0.0001).

## Discussion

Despite extensive *in vitro* and *in vivo* investigation of gut microbial species, we still lack a clear understanding of bacterial physiology at the mucosal surface. Our work shows that human colonic organoids represent a suitable model system to bridge mechanistic studies with animal or human data. We employed colonoids for live visualization of *Enterococcus faecalis* colonization dynamics within a physiologically relevant human mucosal environment, thereby revealing both metabolic and biophysical adaptation strategies. By combining human colonoid-derived tissues, live imaging, and Tn-seq screening, we directly observed *Ef* microcolony formation and identified genetic determinants of fitness under the nutrient-limited and physically complex conditions of the mucus. This allowed us to focus on the early colonization and adaptation events that *Ef* undergoes when transitioning from a liquid culture to a native-like intestinal mucus environment. This process could be similar to the one initially employed by *Ef* to colonize the infant gut, where *Enterococcus* and *Escherichia* species are among the early pioneers, thereby forming a favorable niche for subsequent anaerobic species colonization (50). Our findings show that *Ef* strongly prefers the mucosal niche, rapidly attaches and forms robust, biofilm-like colonies, as previously reported in both germ-free mouse models and patient-derived intestinal samples (12, 35). These results deepen our understanding of how a typical commensal exploits host-derived nutrients and stress-sensing pathways to adapt and stably populate on the mucosa, bridging a key gap between traditional *in vitro* biofilm assays and *in vivo* colonization models.

By growing human intestinal tissue on the basal side of the membrane, we were able to observe bacteria–mucus interactions in real time, capturing the entire process from the initial binding of a single bacterial cell to the formation of a biofilm-like multicellular structure. We consistently observed dense, spatially confined *Ef* colonies associated with mucus, either distributed between thick mucus fibers or embedded within the bulk of the gel matrix, which was reshaped during growth. We obtained these results using a simple yet robust Transwell-based microfluidic platform that supports high-resolution live-cell imaging (44). Such dynamic colonization events are typically obscured in animal models and, to our knowledge, this represents the first real-time microscopic observation of bacterial commensal biofilm-like colony formation in native human intestinal mucus.

Our Tn-seq screen further highlighted the predominance of metabolic flexibility and stress-response systems in early mucosal adaptation. Genes involved in ABC/PTS carbohydrate transporters and amino acid biosynthesis (methionine and tryptophan) were essential for growth on mucus, reflecting the reliance on host-released glycans and the scarcity of free nutrients in the mucosal environment. Particularly, since methionine and tryptophan are essential amino acids for *Ef* (51) and are neither abundant in the mucus nor secreted by the host (52), *de novo* synthesis becomes essential for survival in this niche. Also, the two-component systems (CroRS and BsrRS) emerged as critical sensors of envelope stress (53–55), suggesting that *Ef* must rapidly detect and respond to chemical stressors encountered at the mucosa. Collectively, these data portray *Ef* as a metabolically versatile and environmentally responsive opportunist, capable of fine-tuning its physiology to colonize the colonic mucosa.

To better understand the molecular mechanisms underlying the microcolonies observed by microscopy, we compared our Tn-seq results with known biofilm-associated genes. We found that key biofilm-associated factors, including Fsr- or sortase-regulated adhesins (47, 48), while important in abiotic settings and in the *in vivo* gut colonization model (21), were dispensable for early mucosal colonization. This suggests that the demands of colonizing the mucus environment differ significantly from those of classical *in vitro* biofilm models. Additionally, this multicellular lifestyle, which mimics native biofilm-like structures observed in intestinal tissue, may exploit the mucus matrix itself for protection against flow-mediated clearance and in the absence of other species, while also concentrating enzymatic machinery for mucin degradation. Notably, without external stress, such as antibiotics or competing microorganisms, biofilm formation appears not essential for stable colonization in mucus.

One of the factors we identified as essential for early mucus colonization is the glycosyltransferase BgsB. Mutations of *bgsB* abrogated microcolony formation and fitness in our gut organoid model, highlighting the importance of cell membrane composition and the associated glycolipids for the attachment and growth of *Ef* in clusters within mucus. Future studies will investigate how BgsB-mediated modifications influence host–microbe interactions and whether targeting this pathway could selectively prevent *Ef*’s pathogenic overgrowth. By sequencing the genetic library and imaging colony growth over short periods, we were able to avoid the point at which *Ef* would typically overgrow and infect the tissue. However, the underlying mechanisms that drive the transition of *Ef* from a commensal to a pathogen in the gut, shifting from homeostasis to dysbiosis, remain unresolved. Identifying the factors that influence this transition will be an important next step for this model, which could also facilitate direct drug testing and the examination of polymicrobial communities within the same experimental setup.

Organoids are now becoming indispensable tools for dissecting host–microbe interactions in conditions that most resemble the human mucosal surface, offering an intermediate complexity between oversimplified *in vitro* assays and overly complex and not always relevant animal models. Human colonoids recapitulate key features of the large intestine, such as polarized epithelium, mucin-secreting goblet cells and a native mucus layer. These elements are absent in conventional carcinoma-derived cell lines, yet they are critical for shaping the native-like host environment and investigating microbial colonization in this context. One critical future direction is to develop organoid systems that are capable of generating double-layered mucus (56) and an anaerobic environment (57), which better mimics the native colonic barrier and enables more accurate modeling of host-microbe interactions. In this study, we showed that colonoids uniquely capture the dual biophysical-metabolic interface of the gut mucus niche, making them particularly well suited for studying commensal stability, dysbiosis, and pathogen emergence within the microbiome (58).

## Materials and Methods

### Bacterial culture conditions

Bacteria were cultured in BHI medium (Sigma Aldrich cat. no. 53286-500G) with all liquid cultures incubated at 37 °C without shaking unless otherwise indicated. Strains used can be found in **Table 3**. Antibiotics were used with overnight cultures of fluorescent strains. Spectinomycin (Chemie Brunschwig cat. no. J61820-06) was dissolved in water at 60 mg/ml and stored at −20 °C until use at a working concentration of 120 µg/ml.

### Generation of *E. faecalis* OG1RF Δ*bsrRS* and Δ*croRS* Mutants

Deletion of the *bsrRS* operon (originally annotated as *yclRK*, but renamed here to *bsrRS* for consistency with nomenclature in *Enterococcus faecium* (55)) in the *E. faecalis* OG1RF background was carried out using the temperature-sensitive Gram-positive plasmid pGCP213 (59). To construct the deletion cassette, approximately 0.5 kb regions flanking the *bsrRS* coding sequence were PCR-amplified using primers listed in **Table 1**. Restriction sites for *SphI* and *EcoRI* were incorporated to facilitate cloning. The amplicons included the first and last ∼15 amino acid codons of the gene to minimize unanticipated polar effects. These flanking regions were fused via overlap extension PCR to generate the final insert. Both the insert and pGCP213 plasmid were digested with *SphI* and *EcoRI* (New England Biolabs, USA) and ligated using T4 DNA ligase, following the manufacturer’s protocol (New England Biolabs, USA). The resulting plasmid, pGCP-yclRK, was transformed into competent *E. coli* Stellar cells and verified by PCR and Sanger sequencing. The construct was then introduced into electrocompetent *E. faecalis* OG1RF cells via electroporation, and mutant strains were isolated as previously described (59). Gene deletion was confirmed by PCR and Sanger sequencing of the flanking regions in the mutant strain.

**Table 1.**
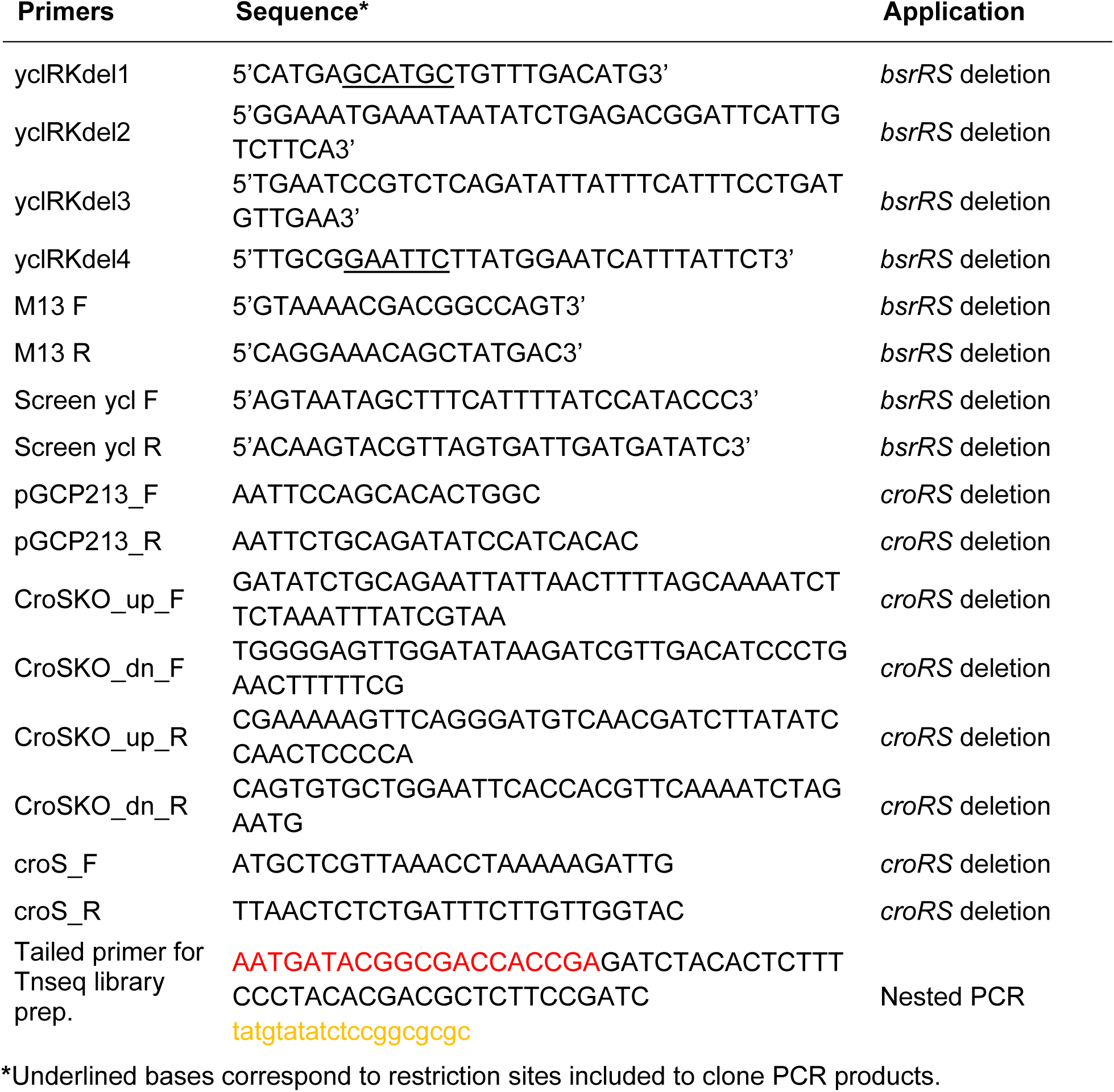
Primers.

Generation of the *croRS* knock-out mutant in *E. faecalis* OG1RF was also performed via allelic replacement using pGCP213, as described previously (59). The pGCP213 plasmid was first linearized by inverse PCR using primers pGCP213_F and pGCP213_R. Approximately 800 bp regions upstream and downstream of the *croS* gene were amplified from the OG1RF chromosome using primer pairs CroSKO_up_F/CroSKO_up_R and CroSKO_down_F/CroSKO_down_R, respectively. These flanking regions were then fused via overlap extension PCR using primers CroSKO_up_F and CroSKO_down_R. The linearized vector and fused insert were ligated using the In-Fusion HD Cloning Kit (Clontech, Takara, Japan) and transformed into *E. coli* Stellar competent cells. Correct constructs were confirmed by PCR and Sanger sequencing, and plasmids were extracted and introduced into *E. faecalis* OG1RF via electroporation.

Transformants were selected on erythromycin-containing media at 30 °C, and chromosomal integrants were selected by growth at 42 °C in the presence of erythromycin. To excise the integrated plasmid via homologous recombination, cultures were serially passaged at 37 °C without antibiotic selection. Erythromycin-sensitive colonies were screened by PCR using primer pair croS_F/croS_R to confirm loss of the *croS* gene. Complete deletion of the *croS* gene, which overlaps with the downstream region of *croR*, was confirmed by whole-genome sequencing. Loss of expression of both CroR and CroS was validated by immunoblotting.

### Transformation of pP_23_::tdTomato into *E. faecalis* OG1RF *Δbph* and *bgsB::Tn*

Transformation procedures were previously described (60). In brief, 1 mL of overnight culture *E. faecalis* in 5 mL M17 medium (HIMEDIA cat. no. M1029-100G) was inoculated in 100 mL fresh SGM17 (M17 broth with 0.5M sucrose (HUBERLAB cat. no. A2211.1000) and 8% glycine (Fisher Scientific cat. no. BP381-1)) and incubated at 37 °C for 18 h. Cells were pelleted at 1,000 g for 10 min at 4 °C. After discarding the supernatant, the pellet was washed with 2 mL ice-cold electroporation buffer (0.5M sucrose with 10% glycerol (Sigma-Aldrich cat. no. G7893-1L), pH7.0). Cells were pelleted at 1,000 g for 10 min at 4 °C. After repeating three washes, electrocompetent cells were resuspended in the electroporation buffer, divided into aliquots, and stored at −80 °C.

To perform electroporation, 1 µL of pP_23_::tdTomato DNA (1.0 µg/µL) was added to thawed electrocompetent cells. The cells were electroporated with 2.5-kV pulse, 25 µF capacitance, and 200 Ω resistance. Cells were recovered at 37 °C for 2 h without aeration and plated on BHI agar plates.

### Culturing human colon organoids

Human colon organoids (donor #1, SCC357, 3d-GRO Merck human distal colon organoids; donor #2, SCC346, 3d-GRO Merck human distal colon organoids) were embedded in Matrigel (Corning cat. no. 356231) and grown into colonoids in a humidified incubator at 37 °C and 5% CO_2_, using human intestinal Start medium. Grown colonoids were passaged as described previously (61). Briefly, Matrigel domes were dissolved in ice-cold Basal medium, collected, and pelleted for 5 min at 200 g, 4 °C. The organoid pellet was resuspended by pipetting in 1 mL of ice-cold Basal medium and an additional 9 mL of ice-cold Basal medium was added. After a second centrifugation step, when no more Matrigel was visible, colonoids were resuspended in freshly thawed Matrigel. A droplet containing 25 µL of the organoid suspension was added to a well of a 24-well plate (Corning cat. no. 3526) and the Matrigel domes were polymerized for 15 min at 37 °C before 500 µL Start supplemented with 2 µM thiazovivin (Sigma-Aldrich cat. no. SML1045) was added. Organoids were passaged every 4–5 days at a ratio of 1:2 or 1:3 depending on the density. The media composition is listed in **Table 2**.

**Table 2.**
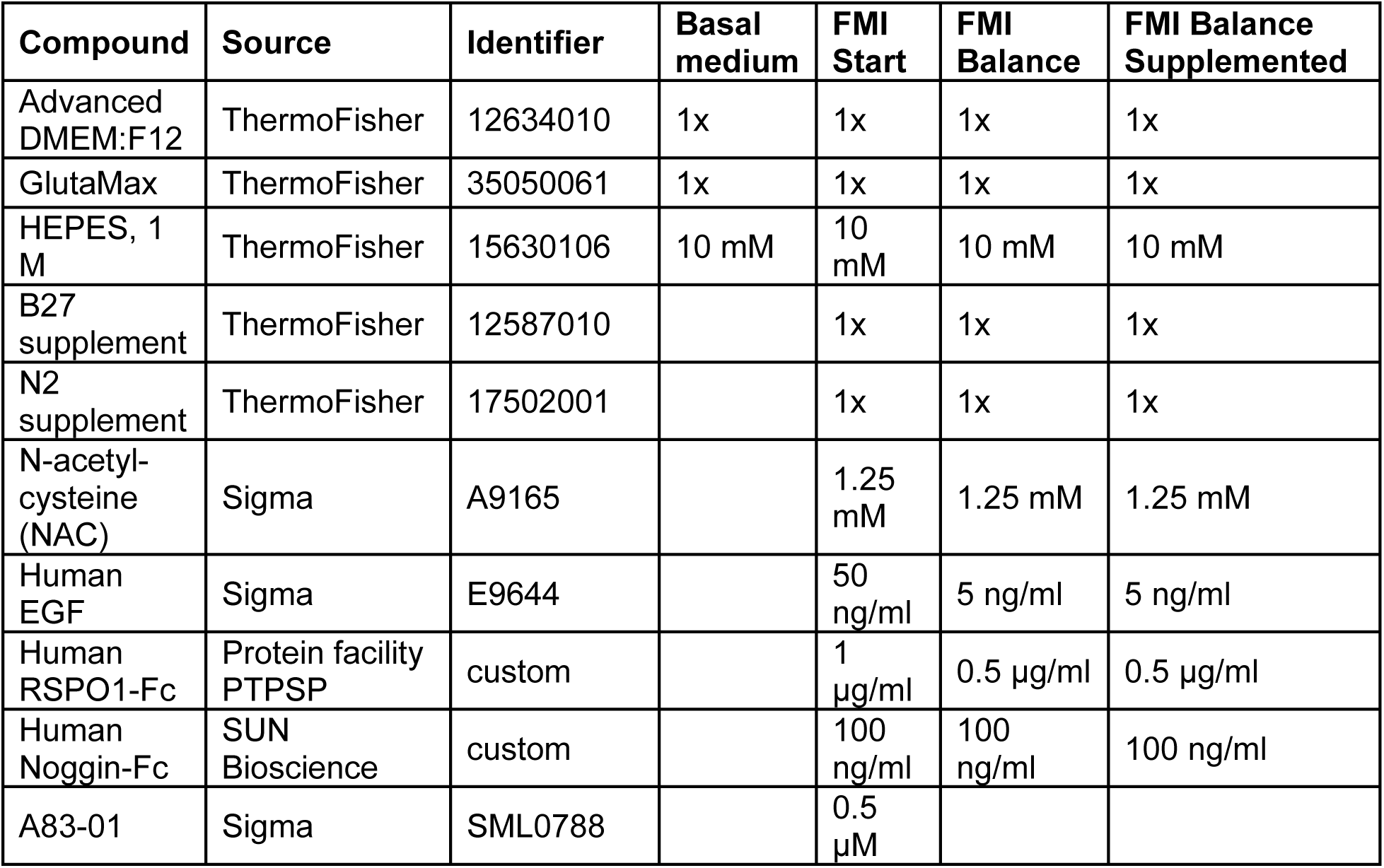
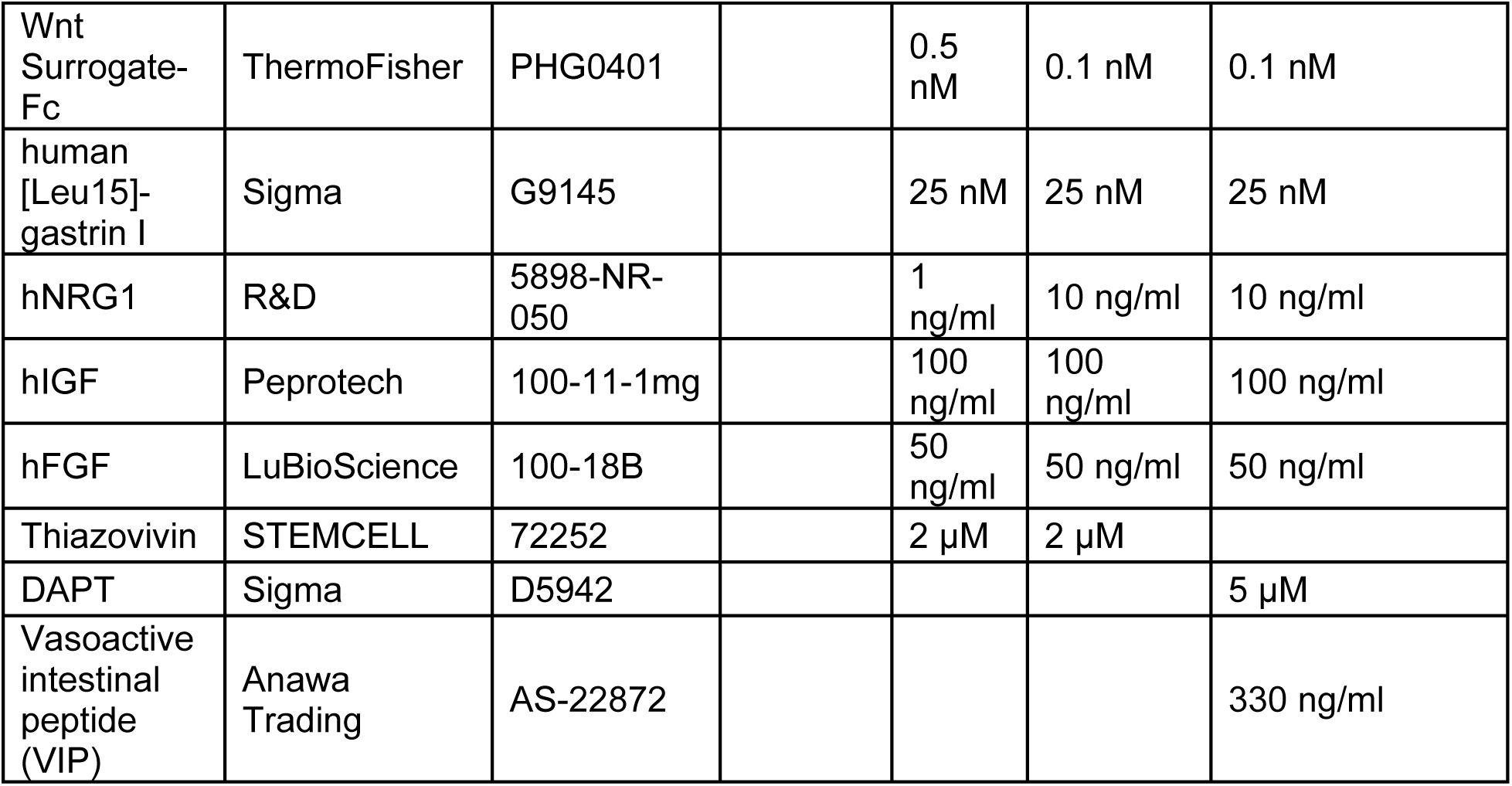
Organoids Media.

**Table 3.** Reagents and strains.

| Reagent or Resource | Source | Identifier |
| --- | --- | --- |
| **Antibodies and lectin staining** |  |  |
| Purified Mouse Anti-Human MUC2 | BD Pharmingen | #555926 |
| Goat anti-Mouse IgG (H+L) Cross-Adsorbed Secondary Antibody, Alexa Fluor 594 | Life Technologies | A11005 |
| Jacalin-biotin | Vector Laboratories | VC-B-1155-M005 |
| Streptavidin-Cy5 | Vector Laboratories | SA-1500-1 |
| **Strains** |  |  |
| OG1RF Tn-seq library | (62) |  |
| OG1RF WT | (63, 64) |  |
| OG1RF WT pDasher-GFP | (43) |  |
| OG1RF *bgsB::Tn* | (46) |  |
| OG1RF $\Delta bph$ | (62) | |
| OG1RF $\Delta bsrRS$ | This study | |
| OG1RF $\Delta croRS$ | This study | |
| OG1RF $\Delta liaFSR$ | (65) | |
| OG1RF *trpX::Tn* | (46) |  |
| OG1RF *metQ::Tn* | (46) |  |
| OG1RF *10303::Tn* | (46) |  |
| OG1RF 11614::*Tn* | (46) |  |
| OG1RF 12205::*Tn* | (46) |  |
| pP<sub>23</sub>::tdTomato | (62) |  |
| OG1RF *bgsB*::*Tn* tdTomato | This study |  |
| OG1RF *Δbph* tdTomato | This study |  |
| **Chemicals** |  |  |
| Brain-heart infusion medium | Sigma-Aldrich | 53286-100G |
| Spectinomycin | Chemie Brunschwig | J61820-06 |
| Rat-tail Collagen I | Corning | 354236 |
| **Materials** |  |  |
| Transwells (upright imaging), 6.5 m, 0.4 μm membrane pores | Sarstedt | 83.3932.041 |
| Transwells (imaging in the chip), 6.5 m, 0.4 μm membrane pores | Corning | 3470 |
| Transwells used for CFUs | Corning HTS | CLS3381 |
| Glass plate for imaging | Cellvis | P12-1.5H-N |

### Culturing differentiated human colon organoids monolayers

To generate differentiate HCO in a Transwell, organoids were collected by centrifugation (5 min, 200 g, 4 °C) and washed until all Matrigel was dissolved as described above and broken down to single cells using 1 mL TrypLE (ThermoFisher cat. no. 12605010) for 7 min in a humidified incubator at 37 °C, 5% CO_2_. Single cells were passed through a 40 µm cell strainer (pluriSelect cat. no. PS-43-10040-40), centrifuged (5 min, 200 g, 4 °C) and resuspended in Start medium, supplemented with 2 μM thiazovivin. Cells were then diluted and seeded onto pre-coated with collagen (50 µg/ml rat tail collagen) Transwell membrane at 100,000 cells in 100 µl to the 24-well Transwells and 45,000 in 50 µl to the 96-well Transwell plate. Five hundred microliter of fresh Balance medium was added to the basal side of the Transwell (200 µl to the 96-well plate wells). In the case of inverted Transwells for high-resolution imaging, Transwells were transferred to a 6-well plate and a custom-made PDMS ring was used to create a well around the basal side of the Transwell insert. 150,000 cells in 100 µl were added to the well in the supplemented Start medium and left to settle for 24 h. The next day, the medium on the apical side of the regular Transwells was changed to the supplemented Balance medium and the inverted Transwells were turned back to a regular configuration with 100 µl of supplemented Balance medium on the apical side and 500 µl on the basal side. Subsequently the medium was changed every other day. When tissues reached full confluency (around 5 days), they were differentiated at the air-liquid interface (ALI) to accumulate mucus with FMI Balance supplemented medium at the basal side. On the third day at ALI tissues were used for experiments with bacterial coculture.

### Bacterial growth at the mucosal surface

All inoculations were performed with bacteria at stationary phase, washed in Hank’s balanced salt solution (HBSS) (ThermoFisher), and diluted to an OD_600_ of 0.5. One microliter of bacterial suspension (24-well) / 0.4 µl (96-well) / 5 µl (inverted Transwell) in HBSS was carefully added to the mucus-filled side and left to settle in the cell culture incubator for at least 30 min until use.

### Tn-seq for growth at the mucosal surface

The transposon library stock was diluted in BHI to an OD_600_ of 0.25 (3 ml) and grown until the deep stationary phase for 14 h. Starting this culture with a high OD_600_ was important to limit the number of generations and consequent pre-selection before the experiment started (∼4 generations for the bulk library), keeping a high transposon saturation in the inoculum. Then, cells were spun down, washed in HBSS buffer, resuspended in HBSS at an OD_600_ of 0.5, and used as inoculum of the Transwell and at OD_600_ of 0.05 as inoculum of the liquid cultures (BHI, Cell Medium (here Basal Medium)). Six different 24-well Transwell inserts for donor SCC357 were infected using 1 µl, which is around ∼250,000 *Ef* cells per insert. The infection progressed for 6 h, which resulted in ∼6 generations without tissue damage by the library. Stationary-phase cells were used so that *Ef* are forced to adapt and grow exclusively from nutrients found in the mucosal environment. After infection, the tissue was homogenized using 100 µl of Triton X-100 (0.1% in PBS). Each insert was scraped until all the tissue was removed. Then, samples from all the inserts were pooled, vortexed and pipetted vigorously, spun down (2 min, 8,500 rpm), washed in HBSS to remove traces of Triton X-100 and spun down again, and the pellets were stored at −80 °C until subsequent processing.

In parallel, we prepared BHI and Cell Medium control samples, in which the inoculum used during the infections was grown in liquids for the same amount of duplication as the infections. For these, the inoculum was diluted in BHI and Basal Medium (3 ml) at an OD_600_ of 0.02 for 3.5 and 5 hours respectively. One milliliter of the culture was spun down, resuspended in HBSS, spun down, washed, and then stored at −80 °C until subsequent processing.

The frozen pellets were processed using the QIAamp DNA Mini kit (Qiagen). First, samples were thawed on ice for 20 min. Bacterial pellets were then resuspended in 200 µl ATL kit buffer + 30 µl 50 mg/ml lysozyme + 20 µl Proteinase K, mixed thoroughly by vortexing, and left shaking at 37°C for 30 min. After that, the steps in the kit were followed exactly. Genomic DNA concentration was measured with Nanodrop and Qubit. Library preparation and sequencing were performed at the Lausanne Genomic Technologies Facility, located at the University of Lausanne.

### Colony forming units

To calculate colony forming units, liquid cultures were serially diluted in HBSS and plated on BHI plates. Transwell cultures were first resuspended in 0.1% Triton X-100 in HBSS (24-well Transwells in 100 µl and 96-well Transwells in 50 µl) as described above, spun down, washed and serially diluted in HBSS. The next day colonies were calculated. Colony forming units/ml were recorded and doublings were calculated as log_2_ (final CFU per ml / initial CFU per ml).

### Confocal imaging of bacterial growth

For labeling mucus and MUC2 immunostaining (Figure 1B), mucus was first labeled with Jacalin-biotin (5 mg/ml) with Streptavidin-Cy5 (40 µg/ml in HBSS). The tissue was fixed with 4% paraformaldehyde in PBS for 30 min at room temperature at the apical side of the Transwell. The fixed tissue was washed three times with PBS, permeabilized with 0.2% Triton X-100 in PBS for 30 min at room temperature, and blocked with the blocking buffer (5% BSA and 0.01% Triton X-100 in PBS) overnight at 4 °C. The tissue was then probed with the anti-MUC2 antibody in the staining buffer (2% BSA in PBS) overnight at 4 °C. After washing three times with PBS, the Goat anti-Mouse IgG-Alexa Fluor 594 secondary antibody in the staining buffer was added to the apical side for one-hour incubation at room temperature. Labeled tissue was washed three times in PBS and stained with 300 nM DAPI for 10 min, followed by three PBS washes.

Imaging bacteria growing in mucus (Figure 1D**, 1E, 4B, 4D, and 4E**) was done as follows. *Ef* strains with plasmids encoding a fluorescent protein (WT expressing Dasher-GFP and *bgsB::*, *Δbph* expressing tdTomato) were grown in BHI broth (containing spectinomycin 120 µg/ml) from a single colony on a BHI plate with 120 µg/ml spectinomycin. Then 500 µl of cells were spun down (2 min at 8000 rpm), washed in HBSS twice, and resuspended in HBSS to an OD_600_ of 0.5. One microliter of the inoculum was used for a regular Transwell and 5 µl for the inverted (most of the unattached bacteria are then washed away in the chip). A 50:50 mixture was applied to the cases where WT and the mutant were grown together (Figure 4D and E).

Epithelium labeling was achieved by incubating the Transwells with 5 µg/ml of CellMask Plasma Membrane Stain Deep Red diluted in the supplemented Balance Medium for 30 minutes in the cell culture incubator at 37 °C. The unbound stain was washed with fresh supplemented Balance Medium. Mucus labeling was done using biotinylated Jacalin (5 mg/ml). A 500 µg/ml Jacalin solution in HBSS was applied at 1 µl on top of the Transwell or at 5 µl to the inverted Transwell. To label Jacalin-biotin, we applied 40 µg/ml of Streptavidin-Cy5 solution in HBSS diluted bacterial culture directly at the infection step (Figure 2 **and** Figure 4).

Confocal imaging was performed using Nikon Eclipse Ti2–E inverted microscope coupled with a Yokogawa CSU W2 confocal spinning disk unit and equipped with a Prime 95B sCMOS camera (Photometrics). Transwell inserts in the upright position (Figure 1 **and** Figure 4) were placed in custom-designed PDMS inserts, plasma-bonded to the glass in 12-well glass bottom plates. Transwells were then imaged using a ×20 water immersion objective (Nikon), 0.95NA, with Di01-T405/488/532/647-filters, 488, 561 and 640 nm laser lines and 100 ms exposure of the fluorescent channels. Brightfield, GFP, Cy3 and Cy5 channels were collected. Images were collected as a z-stack of 2 or 3 µm steps. Fiji was used for the display and the analysis of images. To analyze the volume, image stacks were pre-processed using “Subtract background” function and “Gaussian Blur” filter on the bacteria channel, set at 1 µm and then processed using 3D Objects counter plugin.

To image bacteria-mucus interaction at high resolution under dynamic flow conditions, inverted Transwell inserts were carefully inserted into the microfluidic chip as previously described (44). The chip featured three identical micro-milled open well channels with a glued glass coverslip (ref. 10812, ibidi GmbH, Martinsried, Germany) on the bottom. Metallic needles obtained from Luer-lock connectors (IP721-90; GONANO Dosiertechnik GmbH, Breitstetten, Austria) were then securely attached to the channel inlet and outlet ports. These needles were subsequently connected via Tygon-tube connectors (070534-08L-ND; ID 0.76 mm, wall 0.85 mm, Idex Health & Science GmbH, Wertheim, Germany) to a polytetrafluoroethylene (PTFE) tubing (S1810-08; ID 0.5 mm, OD 1.0 mm, Bohlender GmbH, Grünsfeld, Germany). Additionally, a custom-designed 3D printed lid with holes for the PTFE tubing by Multi Jet Modeling on a Connex 500 printer (Objet) using VeroClear resin at the Additive Manufacturing Workshop (AFA) at EPFL was positioned onto the Transwell inserts (basal compartment) to minimize evaporation of medium and supply fresh medium to the cells. Following the insertion, the microfluidic channels (apical compartment) were manually filled with a 5 ml Luer-lock syringe (Becton Dickinson cat. no. 309649) connected to the PTFE tubing by a metallic needle (GGA725050; GONANO Dosiertechnik GmbH, Breitstetten, Austria). One-milliliter Luer-lock syringes (Becton Dickinson cat. no. 309628) with organoids medium were connected in a similar way through the lid. The syringes were operated via an external syringe pump system (Legato210, KD Scientific, cat. no. 78-8210), perfusing the apical compartment with supplemented SILAC Advanced DMEM/F-12 Flex Media (ThermoFisher) (147.5 mg/l L-arginine hydrochloride (Apollo Scientific), 91.25 mg/l L-lysine hydrochloride (Apollo Scientific), GlutaMAX (ThermoFisher), 10 mM HEPES (ThermoFisher), and MEM NEAA solution (ThermoFisher)) at a flow rate of 5 µl/min and replenishing the basal compartment with organoid medium at 0.5 µl/min.

The chip was held in place in the stage-top temperature chamber (Okolab) at 37 °C, with constant air/CO_2_ mix supply to support cell physiology. Transwells were then imaged using a ×40 water immersion objective, 1.15NA. Timelapses were recorded for > 18 hours, every 30 min. To maintain the water for the objective, a custom-made water pump (a 30 ml syringe (Becton Dickinson cat. no. 301231) filled with water, with the same tubing connection and syringe pump described above) was used. The time indicated in all images is relative to the selected time point (*t* = 0) to enable tracking of the same position in the image stacks over time. Images were analyzed using Fiji. To calculate the gap size and distribution, we have cropped individual large bacterial colonies and for each image calculated horizontal and vertical line profiles, averaged them, and calculated a moving average across all images.

### Tn-seq library sequencing

Library preparation and sequencing (performed at the Lausanne Genomic Technologies Facility, located at the University of Lausanne) procedures were previously described (28) with slight adjustments. Briefly, Genomic DNA (500 ng) was initially sheared with a Covaris S220 using 450 bp insert settings (50 µL in microTUBES with AFA fiber, Peak incident power: 175, Duty factor: 5%, Cycles per burst: 200, time: 50 sec). Libraries were prepared with the xGen DNA MC UNI Library Prep Kit (IDT, protocol version v2) using xGen UDI-UMI adapters (IDT, 15 µM stock). With these adapters, P5 and P7 sequence are inverted compared to Illumina adapters, allowing transposon sequencing directly from read 1 (P5 side) in a single-end run. The purified ligated product was amplified by PCR with a primer specific for the Illumina P7 sequence (CAAGCAGAAGACGGCATACGA) and a second one specific for the transposon sequence (gcatcaccttcaccctctcc) with a 5’-biotin. PCR was performed with the KAPA HiFi HotStart ReadyMix kit (Roche). Cycling conditions were 98 °C for 45 s, followed by 10 cycles of 98 °C for 15 s, 60 °C for 30 s, and 72 °C for 30 s, and a final extension of 1 min at 72 °C. The library was purified with SPRI beads at a 1X ratio.

The PCR product was captured with pre-equilibrated Dynabeads MyOne Streptavidin T1 (ThermoFischer). At least 1 ml of 1X B&W Buffer (5 mM Tris-HCl pH7.5, 0.5 mM EDTA, 1 M NaCl) was mixed with 25 µl of Dynabeads. After 1 min on a magnet, the supernatant was removed, and Dynabeads were washed with the same volume of 1X B&W Buffer twice. The Dynabeads were resuspended with 50 µl of 2X B&W Buffer (10 mM Tris–HCl pH7.5, 1 mM EDTA, 2 M NaCl). Then, 50 µl of the library was mixed with the washed Dynabeads followed by 30 min incubation at RT on a rotator and placed on a magnet for 2 min. After discarding the supernatant, the Dynabeads were washed three times with 100 µl of 1X B&W Buffer before the final elution in 40 µl H_2_O.

Half of the purified capture was used for the nested PCR with the Illumina P7 sequence (see sequence above) and a tailed primer made of the Illumina P5 sequence (red), the TruSeq read 1 primer binding site (black), and a transposon specific binding sequence (orange) AATGATACGGCGACCACCGAGATCTACACTCTTTCCCTACACGACGCTCTTCCGATCtatgtatatctccggcgcgc (for the colored primers, see Table 1). The HiFi HotStart ReadyMix kit KAPA was used for nested PCR amplification, with the same cycling conditions as above (only 9 cycles in this step). The final library was purified with SPRI beads at a 0.7X ratio. It was quantified with a fluorimetric method (QubIT, Thermo Scientifics) and its size pattern analyzed with a fragment analyzer (Agilent).

Sequencing was performed on an Aviti (Element Biosciences) on a Cloudbreak Freestyle high output flow cell for a 150 cycles single-end sequencing run. Clustering was performed with 1 nM library spiked with PhiX (Element Biosciences). Base calling and demultiplexing were done with bases2fastq version: 1.7.and further processed for transposon insertion analysis.

### Tn-seq data processing and analyses

Library sequencing analysis was performed at the Lausanne Genomic Technologies Facility, located at the University of Lausanne. Reads were trimmed for the Tn IR tag, Illumina adapter and low quality scores using cutadapt (v.4.8 parameters: -g “TATGTATATCTCCGGCGCGCCGCGACGCCATCTATGTGTCTAGAGACCGGGGACTTATCAGCCAACCTGTTA;min_overlap=70;max_error_rate=0.1” -A “GATCGGAAGAGCACACGTCTGAACTCCAGTCAC;min_overlap=10” -q 20 --pair-filter first). Only paired-reads with a length larger than 50 nt for read 1 and 30 nt for read 2 were aligned against Enterococcus_faecalis strain OG1RF (Accession number: CP002621.1) genome using BWA (v.0.7.18, parameters: mem -v 3 -T 0 -a -M). Alignment files were deduplicated using UMI_tools (v. 0.5.3, parameters: dedup --method=unique --paired --umi-separator=’:’). The number of reads per insertion site was computed using a custom script that processes the genome read alignments from the BAM file (original or deduplicated). This script first filters the alignments using SAMtools (v 1.19) to select reads based on a specified flag (all alignments flag: -F 132; primary alignments flag: -F 388). It determines the exact position of the first base match on the reference genome by considering the read orientation and CIGAR string. The processed positions are subsequently analyzed and adjusted for any end clipping in the CIGAR string. Finally, it counts the occurrences of each insert site position in the genome and reports them in a sorted wiggle format file. In parallel, the number of read counts per gene locus was summarized with featureCounts (v.1.6.0, parameters: -p -s 0 -M --fraction).

Finally, primary and deduplicated sequences were analyzed in Transit with the Tn5 ‘resampling’ method within TRANSIT (GUI mode) to assess the conditional essentiality of genes between two conditions in three distinct comparisons. Specifically, we tested ‘experimental sample’ in comparison to the second as the ‘control sample’: (1) mucus growth (5_muc) versus BHI control (3_bhi), (2) mucus growth versus cell medium control (4_dmem), (3) cell medium growth versus BHI control. The initial library comparison to the inoculum is also available and shows minimal differences (1_start and 2_inn) (see **Table S1**).

### Enrichment analyses of functional processes using KEGG pathways and GO terms

For the classification of gene functions, a custom annotation (see **Table S2**, based on the previous version in (46)) and the Kyoto Encyclopedia of Genes and Genomes (KEGG) database for the OG1RF strain were used. We selected 333 genes as significant (having log2FC < −2 in comparison to both the BHI and the cell medium). Gene ontology terms based on “GO_process” were applied from the NC_017316.1 full genome sequencing annotation in the GenBank and grouped accordingly using a custom-written R code, and the number of genes in each category was plotted in Figure 3C.

### Growth curve measurement

Overnight cultured *Ef* WT and mutants in BHI medium were spun down at 8,000 rpm for 2 min. The pellets were washed twice with PBS and diluted to OD_600_ = 0.02 in 100 µl of BHI medium or minimal medium described above, with 1% (W/V) of D-galactose (Sigma-Aldrich), GalNAc (Sigma-Aldrich), GlcNAc (Adipogen), D-glucose (Carl ROTH), or D-mannose (Sigma-Aldrich) and loaded to a 96-well plate. The plate was sealed with parafilm and subjected to the SPECTROstar Nano plate reader (BMG LABTECH) for measuring OD_600_ every 10 min without shaking at 37 °C.

### Statistical analysis

All statistical tests were run in R using rstatix::t_test toolbox for a Welch’s t-test, which doesn’t assume equal variances between groups. Pairwise comparisons were performed between WT and each mutant group individually. We adjusted the p-values to be displayed as: p < 0.0001 ∼ “****”, p < 0.001 ∼ “***”, p < 0.01 ∼ “**”, p < 0.05 ∼ “*” and “ns” for p >= 0.05.

## Acknowledgments

We thank the Lausanne Genomic Technologies Facility (GTF) for their sequencing services, the EPFL Protein Production and Structure Core Facility, particularly Kelvin Lau and Florence Pojer, for providing the organoid growth factors, EPFL Additive Manufacturing Workshop (AFA) for 3D printing, Kline and Willet labs for providing the *Ef* transposon library and mutants. We also thank Zaïnebe Al-Mayyah for laboratory technical assistance, and members of the Persat, Hierlemann and Kline labs for constructive feedback throughout the development of the project. We also thank Prisca Liberali and Koen Oost for their insights on human colon organoid protocols. This work was supported by the Swiss National Science Foundation (SNSF) Projects grant 310030_204190. S.M. is the recipient of a Peter and Traudl Engelhorn Postdoctoral Fellowship (2022–2024). A.K., J.A.B., A.H. and A.P. are members of the NCCR AntiResist. Authors declare that they have no competing interests.

## Author contributions

Conceptualization: S.M., A.P. Data curation: S.M., P.H. Formal Analysis: S.M., P.H.

Funding acquisition: S.M., A.P. Investigation: S.M., P.H.

Methodology: S.M., A.K., C.M.A., C.C.W., J.A.B., P.Y.C., J.L.E.W.

Project administration: S.M., P.H., A.P. Supervision: A.P., K.K., A.H. Visualization: S.M., P.H.

Writing – original draft: S.M., P.H., A.P.

Writing – review & editing: P.H., S.M., C.M.A., C.C.W., K.K., A.K., A.P.

