## Supplementary material for "Adaptation of *Enterococcus faecalis* to intestinal mucus revealed by a human colonic organoid model": Figure S1-4

### Supplementary Figures

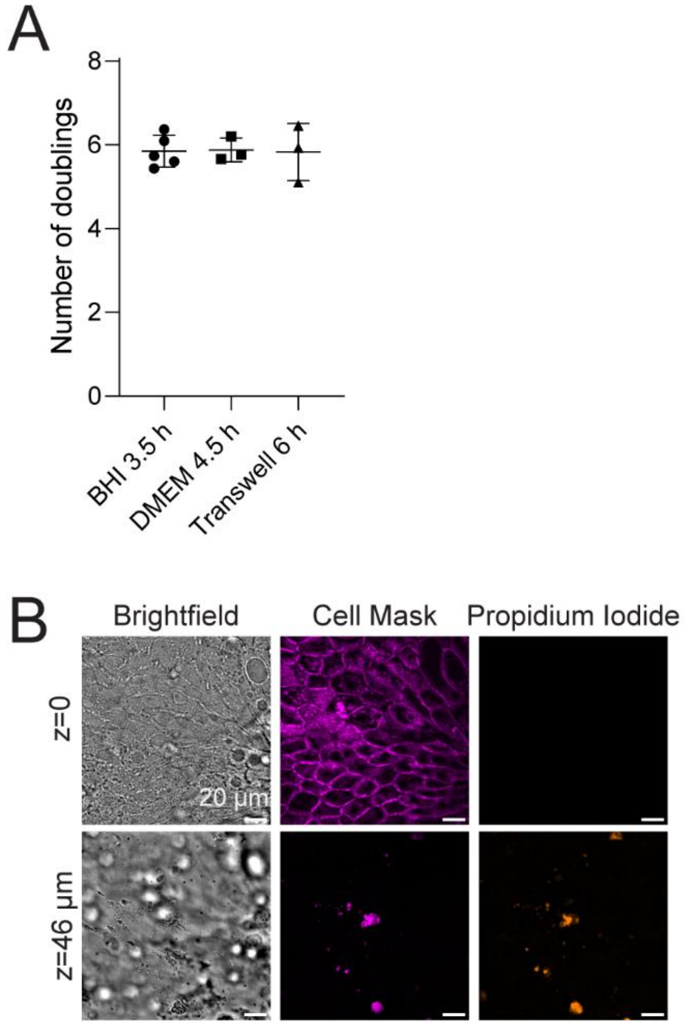


**Figure S1. Calibration of the Tn-seq condition.** (A) Incubation times of the library were calibrated to obtain similar doublings in rich medium (BHI), colonoid medium (DMEM), or Transwells. Each datapoint is a biological replicate. Mean and standard deviation are shown. (B) Epithelial cells do not show a significant loss of integrity after growing the library in mucus. The epithelial layer is stained with CellMask-DeepRed (magenta), and dead cells are labeled with propidium iodide (orange). Dead cells are only present in the mucus layer as in typical organoid cultures.

A

Tryptophan ABC-family transporters

| Locus tag | Name | Mucus / BHI Medium | Mucus / Cell Medium |
|---|---|---|---|
| OG1RF_RS05315 (OG1RF_11022) | *trpX* | -4.77 | -2.95 |
| OG1RF_RS05320 (OG1RF_11023) | | -4.74 | -2.97 |
| OG1RF_RS05325 (OG1RF_11024) | | -2.86 | -1.59 |

Methionine ABC-family transporters

| | | | |
|---|---|---|---|
| OG1RF_RS09800 (OG1RF_11916) | *metQ* | -3.38 | -2.59 |
| OG1RF_RS09805 (OG1RF_11917) | *metI* | -1.78 | -0.99 |
| OG1RF_RS09810 (OG1RF_11918) | *metN* | -2.14 | -1.24 |

PTS-family sugar transporters

| | | | |
|---|---|---|---|
| OG1RF_RS08280 (OG1RF_11613) | | -1.35 | -0.99 |
| OG1RF_RS08285 (OG1RF_11614) | | -2.03 | -1.38 |
| OG1RF_RS08295 (OG1RF_11616) | | -1.4 | -0.83 |

ABC-family sugar transporters

| | | | |
|---|---|---|---|
| OG1RF_RS01655 (OG1RF_10301) | | -3.63 | -3.15 |
| OG1RF_RS01660 (OG1RF_10302) | | -2.64 | -1.59 |
| OG1RF_RS01665 (OG1RF_10303) | | -3.76 | -3.43 |
| OG1RF_RS11300 (OG1RF_12203) | | -1.48 | -1.15 |
| OG1RF_RS11310 (OG1RF_12205) | | -3.76 | -3.4 |
| OG1RF_RS11315 (OG1RF_12206) | | -1.54 | -1.28 |
| OG1RF_RS11320 (OG1RF_12207) | | -1.29 | -1.0 |

log2FC in fitness: 0, -1, -2, -3, -4, -5

B

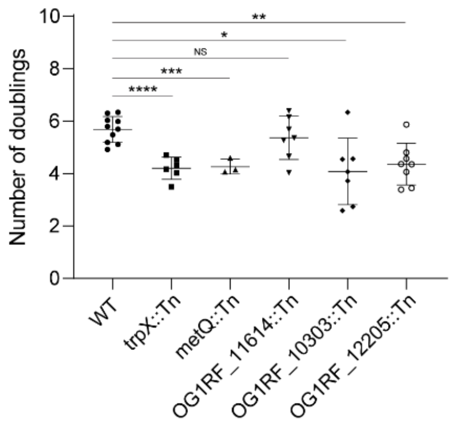


C

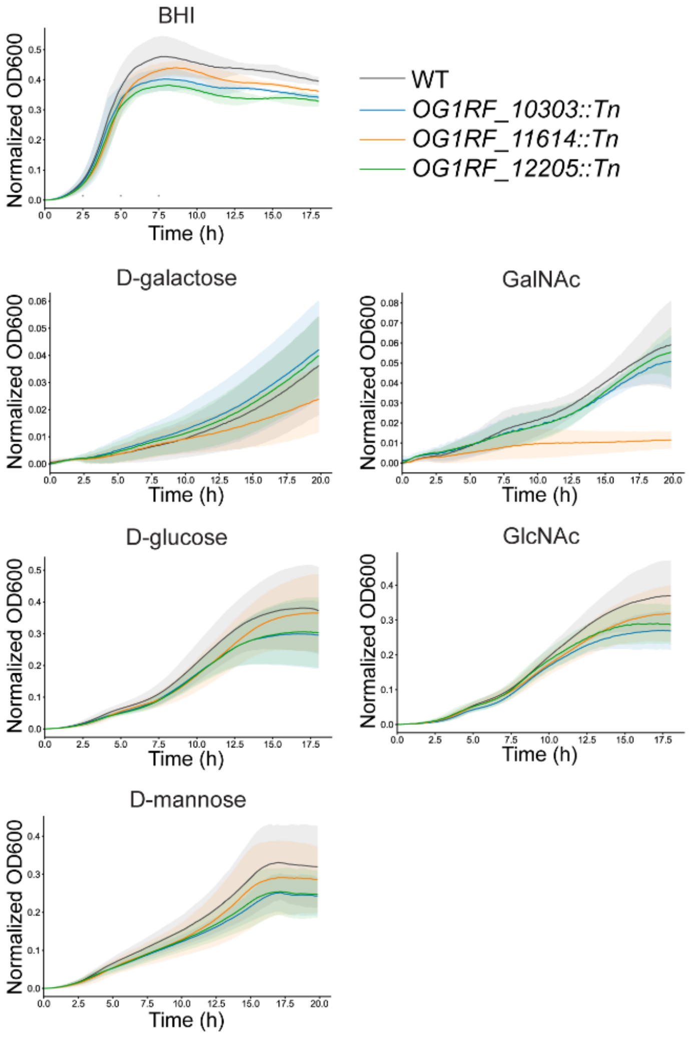


**Figure S2. The PTS and ABC sugar and amino acid transporting systems are required for *Ef* persistence in colonic mucus.** (A) Table of log2FC values calculated from the Tn-seq experiment for each selected gene among ABC- and PTS-transporters. (B) Colony forming units (CFU/ml) quantification of the mutants grown in mucus in comparison with the wild type *E. faecalis (****, p< 0.0001; ***, p<0.001, **, p<0.01, *, p<0.05, NS, p > 0.05).* Mean and standard deviation are shown. (C) Growth curves of transposon mutants and WT grown in BHI or liquid minimal medium supplemented with 1% indicated sugar in a 96-well plastic plate. Mean (line curves) and standard deviation (shade with corresponding color) are shown (n = 3).

A

Two-component regulatory systems

| Locus tag | Name | Mucus / BHI Medium | Mucus / Cell Medium |
|---|---|---|---|
| OG1RF_RS05350 (OG1RF_11029) | *bsrR* | -1.53 | -0.93 |
| OG1RF_RS05355 (OG1RF_11030) | *bsrS* | -1.33 | -0.83 |
| OG1RF_RS12980 (OG1RF_12535) | *croR* | -1.74 | -1.1 |
| OG1RF_RS12985 (OG1RF_12535) | *croS* | -2.19 | -0.72 |
| OG1RF_RS11340 (OG1RF_12211) | *liaR* | -3.37 | -2.9 |
| OG1RF_RS11345 (OG1RF_12212) | *liaS* | -2.57 | -2.04 |

log2FC in fitness: 0, -1, -2, -3, -4, -5

B

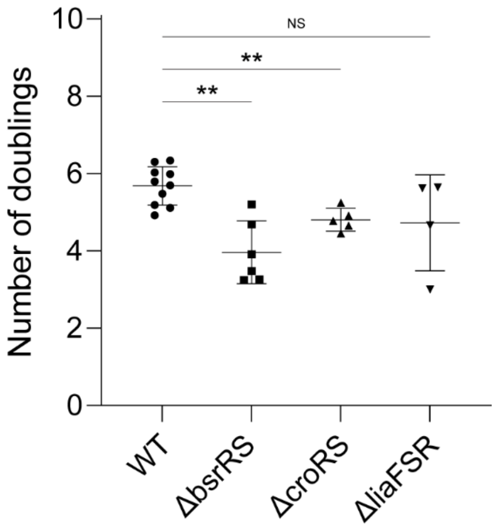


**Figure S3. The two-component regulatory systems are required for *Ef* persistence in colonic mucus.** (A) Table of log2FC values calculated from the Tn-seq experiment for each gene in the three selected two-component regulatory systems. (B) Colony forming units (CFU/ml) quantification of the deletion mutants grown in mucus in comparison with the wild type *E. faecalis* *(**, p< 0.01; ns, p > 0.05).*

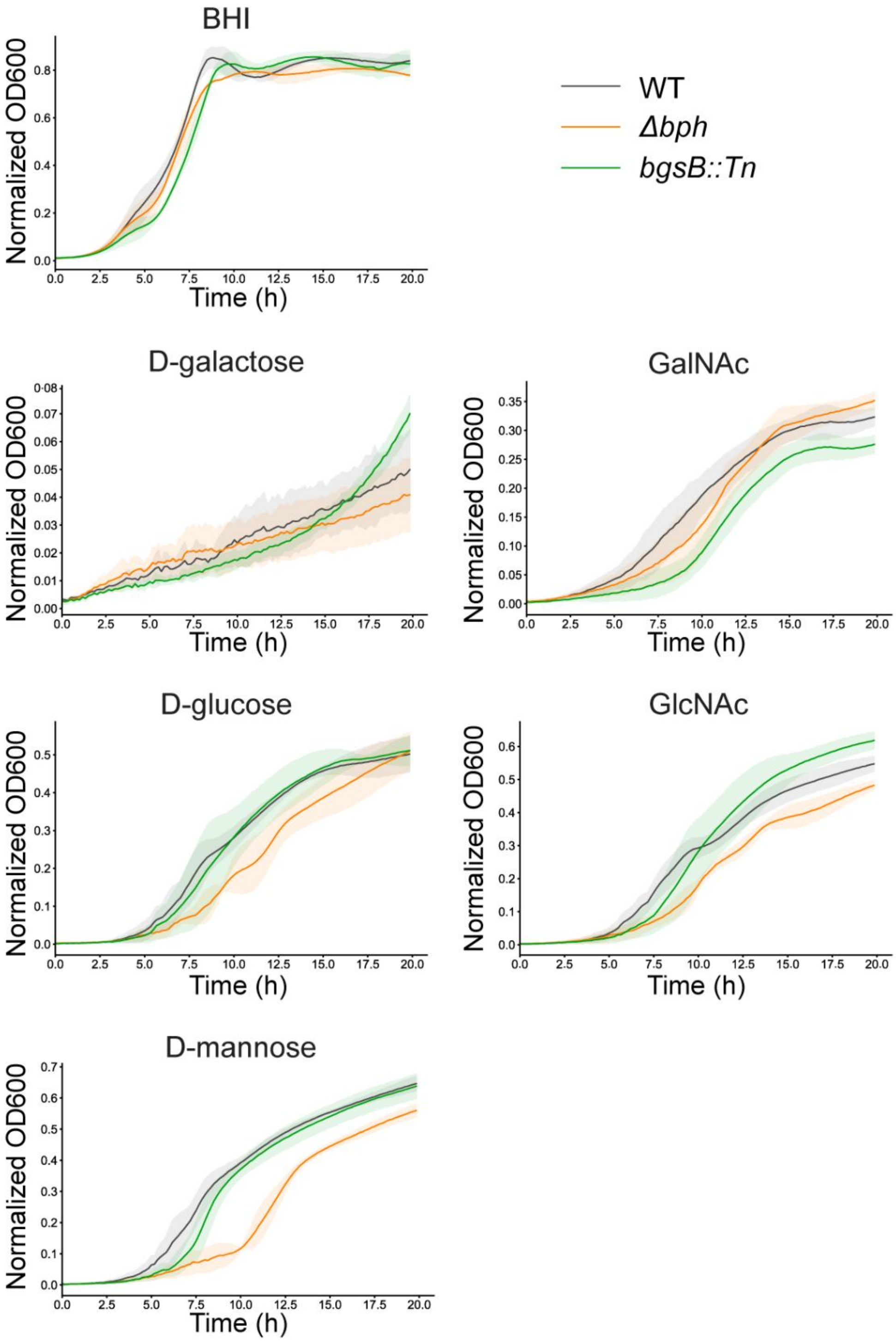


**Figure S4. *Δbph* mutant showed delayed exponential growth in mannose compared to WT.** Growth curves of *Δbph* and *bgsB::Tn* mutants and WT grown in BHI or liquid minimal medium supplemented with 1% indicated sugar in a 96-well plastic plate. Mean (line curves) and standard deviation (shade with corresponding color) are shown (n = 3).
